# Active trachoma cases in the Solomon Islands have varied polymicrobial community structures but do not associate with individual non-chlamydial pathogens of the eye

**DOI:** 10.1101/134213

**Authors:** Robert M R Butcher, Oliver Sokana, Kelvin Jack, Eric Kalae, Leslie Sui, Charles Russell, Joanna Houghton, Christine Palmer, Martin J Holland, Richard T Le Mesurier, Anthony W Solomon, David C W Mabey, Chrissy h. Roberts

## Abstract

**Background:** Several non-chlamydial microbial pathogens are associated with clinical signs of active trachoma in trachoma-endemic communities with a low prevalence of ocular *Chlamydia trachomatis* (*Ct*) infection. In the Solomon Islands, the prevalence of *Ct* among children is low despite the prevalence of active trachoma being moderate. We therefore set out to investigate whether active trachoma was associated with a common non-chlamydial infection or with a dominant polymicrobial community dysbiosis in the Solomon Islands.

**Methods:** We studied DNA from conjunctival swabs collected from 257 Solomon Islanders with active trachoma and matched controls. Droplet digital PCR was used to test for pathogens suspected to be able to induce follicular conjunctivitis. Polymicrobial community diversity and composition were studied by sequencing of hypervariable regions of the 16S ribosomal ribonucleic acid gene in a subset of 54 cases and 53 controls.

**Results:** Although *Ct* was associated with active trachoma, the number of infections was low (cases: 3.9%, controls: 0.4%). Estimated prevalence (cases, controls) of each non-chlamydial infection was as follows: *S. aureus* (1.9%, 1.9%), *Adenoviridae* (1.2%, 1.2%), coagulase-negative *Staphylococcus* (5.8%, 4.3%), *H. influenzae* (7.4%, 11.7%), *M. catarrhalis* (2.3%, 4.7%) and *S. pneumoniae* (7.0%, 6.2%). There was no statistically significant association between clinical signs of trachoma and presence or load of any of the non-*Ct* infections that were assayed. Inter-individual variations in the conjunctival microbiome were characterised by differences in the levels of *Corynebacterium*, *Proprionibacterium*, *Helicobacter* and *Paracoccus*, but diversity and relative abundance of these specific genera did not differ significantly between cases and controls.

**Discussion:** It is unlikely that the prevalent trachoma-like follicular conjunctivitis in the Solomon Islands has a dominant bacterial aetiology. Before implementing community-wide azithromycin distribution for trachoma, policy makers should consider that clinical signs of trachoma can be observed in the absence of any detectable azithromycin-susceptible organism.

## Introduction

Trachoma, caused by *Chlamydia trachomatis* (*Ct*), is the leading infectious cause of preventable blindness (1), and is targeted for elimination as a public health problem by 2020 through the SAFE strategy (**S**urgery, **A**ntibiotics, **F**acial cleanliness and **E**nvironmental improvement). The decision to implement community-wide trachoma control interventions, which include mass drug administration (MDA) with azithromycin, is based on population prevalence estimates of one clinical sign of active trachoma (trachomatous inflammation-follicular [TF]) in the 1-9-year-old age group (2). In the Solomon Islands, the prevalence of TF is sufficient to warrant MDA but the prevalence of trachomatous trichiasis (TT) suggests trachoma may not pose a significant public health problem. A recent trachoma survey in a treatment-naïve population of the Solomon Islands estimated 26% of 19-year-olds to have TF but, surprisingly, conjunctival *Ct* infection was detected in only 1.3% of that age group (3). Whilst *Ct* infection is not always detectable in TF cases (4, 5), infection prevalence in the Solomon Islands is far lower than is seen in other countries with similar TF prevalence (3).

*Ct* is not the only pathogen to associate with signs of TF. A number of differential diagnoses for follicular conjunctivitis are described (6), some of which are rare and distinguishable by patient history (such as Parinaud’s oculoglandular syndrome). Other causes of conjunctivitis which can present with follicles are *Streptococcus pneumoniae*, *Haemophilus influenzae* (7), and adenovirus (8). Non-ocular *Chlamydia* serotypes (9) and non-*trachomatis Chlamydia* species (10) have also been suggested as possible causes of TF. A number of studies have examined conjunctival microbiology in trachoma-endemic settings. A greater diversity of pathogens can be cultured from the conjunctivae of people with TF compared to counterparts without TF (7). Among bacterial species isolated during a study in Tanzanian children, *Streptococcus pneumoniae*, *Haemophilus influenzae* B and *Haemophilus influenzae* non-type B associated more strongly than *Ct* with clinical signs of active trachoma (7). In The Gambia, *S. pneumoniae* and *H. influenzae* type B infection correlated closely with signs of active trachoma in communities which had received MDA, whereas *Moraxella catarrhalis* and *Staphylococcus aureus* did not (11). Common non-chlamydial pathogens have associated with trachomatous scarring (TS) (12) and recurrence of TT after surgery (13) in some, but not all (14, 15), studies.

Although traditionally thought to be ‘sterile’, several studies have now described polymicrobial communities colonising the conjunctival epithelium (16, 17). *Staphylococcus*, *Streptococcus*, *Haemophilus* and *Moraxella* genera can be readily cultured from swabs taken from the inferior fornix (7, 11, 12). Attempts to detect bacteria from the conjunctiva have shown community diversity and composition to vary significantly between individuals. It is unclear whether a ‘core’ microbiota persists at the conjunctiva, but profiles closely related to that of the skin have been reported (18). *Corynebacterium*, *Propionibacterium*, *Staphylococcus* and *Streptococcus* have been consistently dominant, whereas other genera such as *Acinetobacter*, *Brevundimonas*, *Pseudomonas*, *Bradyrhizobium*, *Sphingomonas*, *Bacillus*, *Simonsiella* and *Elizabethkingia* have been identified more sporadically (19–21). Conjunctival polymicrobial community composition is known to vary with age and season (22) and appears to be responsive to external stimuli such as regular contact lens wear (23). The conjunctival microbiome varies significantly between people with trachomatous scarring and those without, however, significant associations between microbiome and follicular conjunctival inflammation or microbiome and other diseases have yet to be described (22, 24).

We hypothesised that clinical signs of active trachoma in the Solomon Islands where ocular *Ct* is uncommon could be explained by a common non-chlamydial infection or by a dominant polymicrobial community dysbiosis.

## Methods

### Study ethics

A parent or guardian gave written informed consent for each child to take part in the survey in accordance with the Declaration of Helsinki. The protocol was approved by the London School of Hygiene & Tropical Medicine (6319/6360) and the Solomon Islands National Health Research (HRC13/18) Ethics Committees.

### Study population

The samples tested during this study were a sub-set of specimens from a population-based trachoma prevalence survey of 3674 people (1135 children aged 1–9 years) in 32 clusters from Temotu and Rennell & Bellona Provinces, Solomon Islands; data from that survey are presented elsewhere (3). At the time of the survey, no trachoma interventions had been implemented in the Solomon Islands. Sterile swabs were passed three times over the right conjunctiva of every child before storage in 300 μL RNALater^®^ (Life Technologies). Swabs were kept cool in the field for up to 24hrs then frozen (3). Inclusion criteria for cases (n = 257) were age 1–9 years, detectable human DNA in the conjunctival swab and a field grade of TF or trachomatous inflammation—intense (TI) in the right eye. An equal number (n = 257) of age-, gender- and island-matched controls without TF or TI were selected using the ‘e1071’ R package. A random subset of cases (n = 54) and controls (n = 53) were further characterized using V1–V3 16S rRNA gene sequencing. Field controls (clean swabs passed within 20 cm of seated participants, without touching them, and then treated identically to subjects’ specimens), extraction controls (extraction carried out in the absence of a specimen and PCR carried out on the eluate) and PCR controls (PCR carried out in the absence of any added material) were used to determine background levels of microbial contamination in the 16S rRNA gene sequencing protocol.

### Droplet digital PCR

DNA was extracted from conjunctival swabs using the Qiagen AllPrep DNA/RNA kit. We chose not to use mechanical lysis on these low biomass specimens due to the reported lower yields compared to chemical lysis (25). All specimens have previously been tested for *Ct* and *Homo sapiens* Ribonuclease P protein subunit p30 (RPP30, acting as endogenous control) using a validated assay (26); the community prevalence of ocular *Ct* infection is described elsewhere (3).

Duplex ddPCR assays were developed for (1) *Adenoviridae* (27) and *S. pneumoniae* (28), (2) *H. influenzae* (29) and *M. catarrhalis* (30, 31), and (3) *S. aureus* and coagulase-negative *Staphylococcus* (32, 33) based on published assays. Each assay contained 10 μL 2× ddPCR Supermix for probes (Bio-rad, Hemel Hempstead, UK), primers and probes at custom concentrations (Table 1) and 8 μL of template DNA. Thermal cycling was 10’00’’ at 95°C then 40x (0’15’’ at 95°C | 1’00’’ at 60°C) then 12’00’’ at 98°C (26). A positive result for a clinical specimen was defined as >95% confidence of non-zero target load, as described previously (26). Assay performance was assessed by repeat testing of PCR product dilution series’ and cultured pathogen material before rolling out to clinical samples. The reproducibility, linearity and limits of detection of each assay were considered appropriate (Supplementary Table 1) and were also similar to published performance of the *Ct* assay used (26).

**Table 1.**
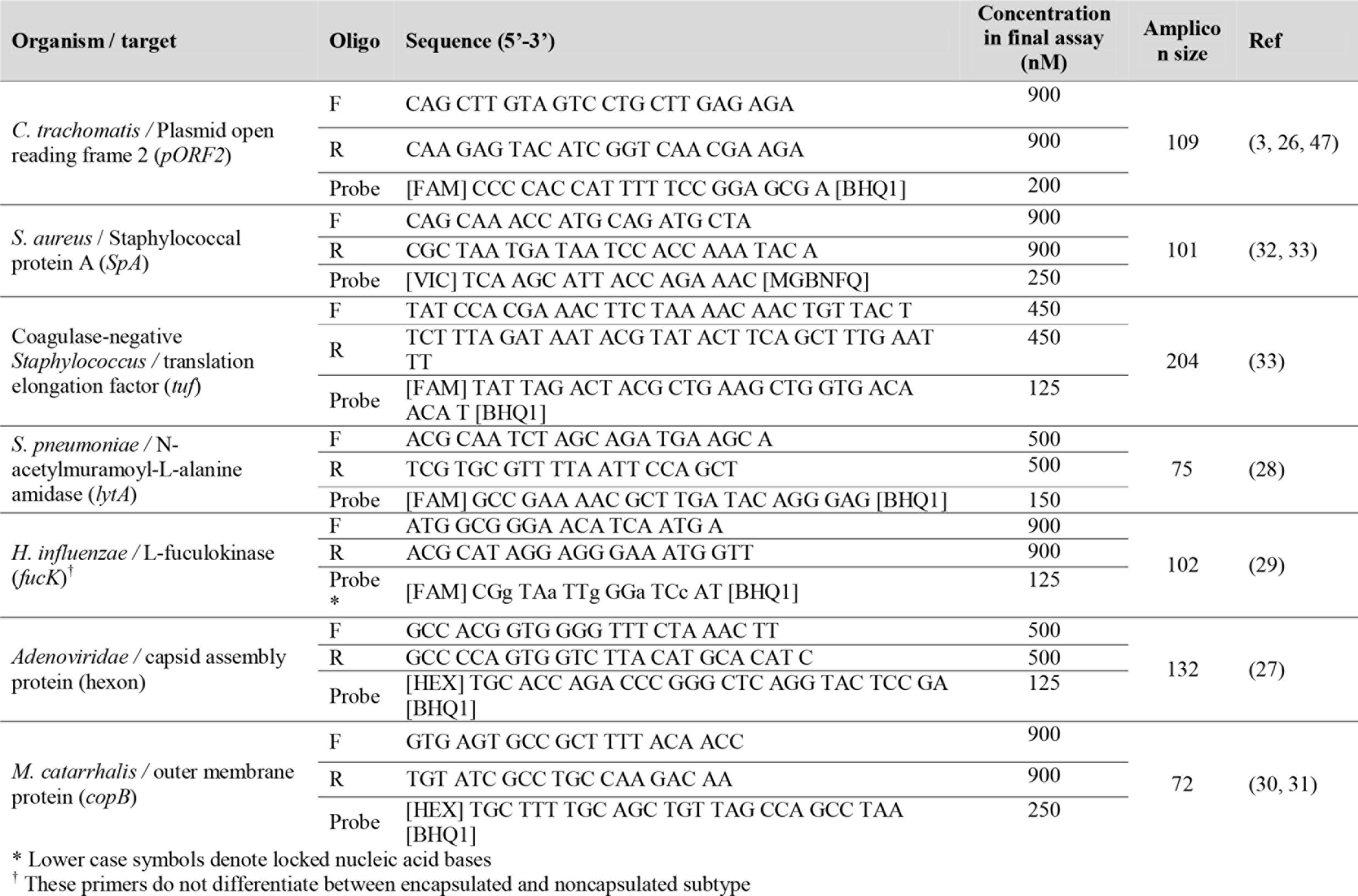
Oligonucleotides used in this study, with assay concentrations derived from 618 *in vitro* optimisation.

### 16S rRNA gene sequencing

An approximately 530-bp region of the 16S ribosomal RNA (rRNA) gene (variable regions 1-3) was amplified using forward (modified 27F; 5’-[adaptor]-AGAGTTTGGATCCTGGCTCAG-3’) and custom barcoded reverse primers (534R; 5’-[adaptor]-[barcode]- AGTCAGTCAGCCATTACCGCGGCTGCTGG-3’). Each 15 μL reaction contained 7.5 μL 2x Phusion High Fidelity Master Mix (New England Biosciences, MA, USA), 0.45 μL DMSO, 0.1 μM primers and 5.55 μL DNA. The thermal cycling conditions were 0’30” at 98°C then 31x (0’10” at 98°C | 0’30” at 62°C | 0’15” at 72°C) then 7’00” at 72°C. Amplicons were cleaned using 0.6 v/v AMPure XP beads (Beckman Coulter, CA, USA), quantified using Qubit (Thermo Fisher Scientific, MA, USA) then pooled at equimolar concentrations. The 4 nM sequencing library was mixed 0.75 v/v with a 4 nM Phi-X control library (Illumina, CA, USA). 3 μL of 100 μM custom read primers were mixed in to wells 12 (Read 1; CTACACTATGGTAATTGTAGAGTTTGGATCCTGGCTCAG), 13 (Index; CCAGCAGCCGCGGTAATGGCTGACTGACT) and 14 (Read 2; AGTCAGTCAGCCATTACCGCGGCTGCTGG) of the MiSeq reagents cartridge before 2 x 300 bp paired-end sequencing with the 600-cycle MiSeq v3 sequencing reagent kit on the MiSeq platform using a standard protocol (Illumina, CA, USA). 96 uniquely barcoded specimens were run in multiplex on each MiSeq run.

### Data analysis

ddPCR data were analysed using R version 3.2.2 (34). Binomial univariate regression was used to test the relationship between each individual infection and active trachoma. The most accurate final multivariate model, determined by lowest Akaike Information Criterion (AIC) value, was determined by step-wise removal of variables from a binomial multivariate regression model incorporating all tested pathogens. The chance of the *Ct* association occurring due to chance was assessed by counting the number of significant results obtained from randomly re-ordering the *Ct* data 1000 times.

Raw 16S amplicon sequences were directly assigned to genera using Illumina BaseSpace ‘16S Metagenomics’ app version 1.0.1.0, which uses algorithms from Wang and colleagues (35). Two clinical specimens yielded fewer than 1000 reads in total and were removed from the analysis. Amplicons were sequenced from six no-template control (NTC) specimens (two ‘field’ controls, two ‘extraction’ controls, two ‘PCR’ controls, defined above) to identify any contaminants endogenous to the collection, PCR or sequencing process. Any genera represented by more than 600 reads in the six combined NTCs (average 100 reads per NTC) were eliminated from the clinical specimen analysis. This resulted in the removal of 21 genera, which accounted for 95.1% of the reads from NTCs and 88.3% of the reads from clinical specimens. The genera removed included common contaminants of reagent kits (36) such as *Pelomonas*, as well as some previously described conjunctival microbiota constituents such as *Brevundimonas*, *Staphylococcus* and *Streptococcus* (19, 22).

Between-group differences in the Shannon Diversity Index (a measure of diversity which increases with increasing species abundance and evenness) and Inverse Simpson Index (another measure of diversity where 1 is infinite diversity and 0 is no diversity) were compared using a t-test. Discriminant Analysis of Principal Components (DAPC), a multivariate clustering method to analyse highly dimensional datasets, was performed using the ‘adegenet’ package in R (37) and cross-validation was used to determine the optimal number of principal components to include in a discriminant function aimed at separating cases from controls on the basis of their polymicrobial community structures.

## Results

### Specimen set demographics

A total of 257 cases and 257 controls (n = 514) were tested. All 257 cases had TF, and one also had TI. Case status was defined by clinical signs in the right eye but respectively 236/257 (91.8%) and 8/257 (3.1%) of the cases and controls had TF in the left eye. Both case and control groups were 38% female. Mean age was 5.6 years (cases) and 5.5 years (controls, student’s t test p = 0.76). There was no significant difference between cases and controls in terms of the clusters represented (Kolmogorov-Smirnov test p = 0.97) and all 32 clusters were represented. The case specimens had higher loads of human DNA than the controls (18030 versus 9354 copies/swab; p = 0.00003). There were no significant differences in age, gender or location within the subset selected for 16S-amplicon sequencing.

### Quantitative PCR tests for ocular pathogens

In this study, 19.8% of children had evidence of infection with at least one of the targeted organisms (Table 2). We considered the prevalence of *Ct* in this sample set to be too low (96.1% of TF cases were *Ct*-infection negative) to account for the level of active disease. The prevalence of *Adenoviridae* (1.2%) and *S. aureus* (1.9%) were both very low. There was no association between active trachoma and infection with *H. influenzae*, *S. pneumoniae*, *S. aureus*, coagulase-negative *Staphylococcus* or M. *catarrhalis. Ct* infection was associated with active trachoma (logistic regression p = 0.026; odds ratio: 10.4). The association between *Ct* and active trachoma was still significant in a multivariate analysis (p = 0.025). The permuted p-value for the association between *Ct* and active trachoma was highly significant (p = 0.001). Active disease was neither associated with infection with at least one pathogen (any pathogen, p = 0.185) nor the number of concurrent infections in the same eye (number of concurrent pathogens, p = 0.207). Infection loads of those who tested positive are shown Figure 1. Despite some numerical differences between groups, none of the differences were statistically significant.

**Table 2.**
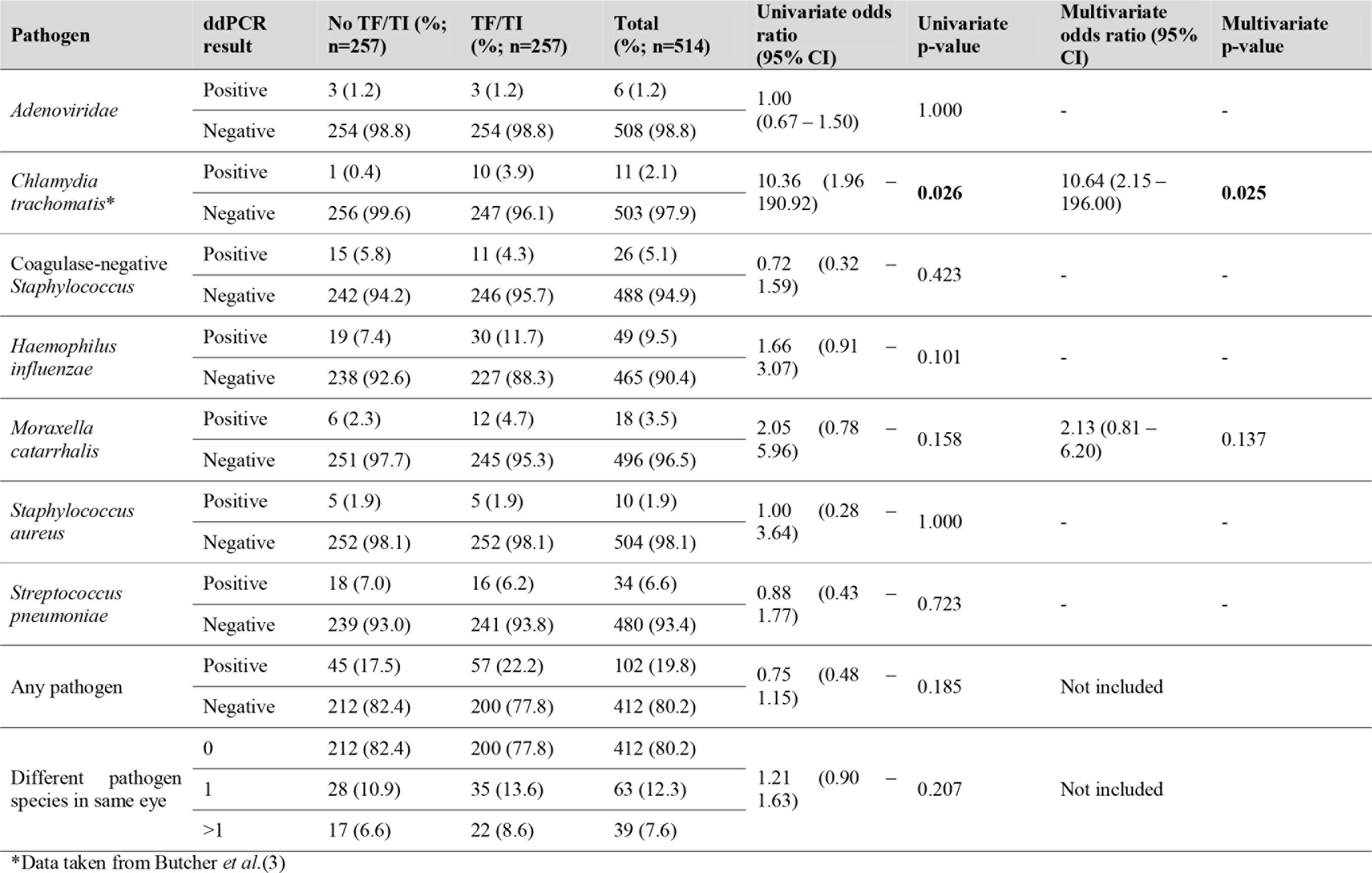
Cases of infection in specimens from children with and without active disease. The relationship of those infections to trachoma has been tested with univariate logistic regression, and then stepwise removal of variables from multivariate regression model was used to determine the final multivariate regression model that provided the best fit for the data.

**Figure 1.**
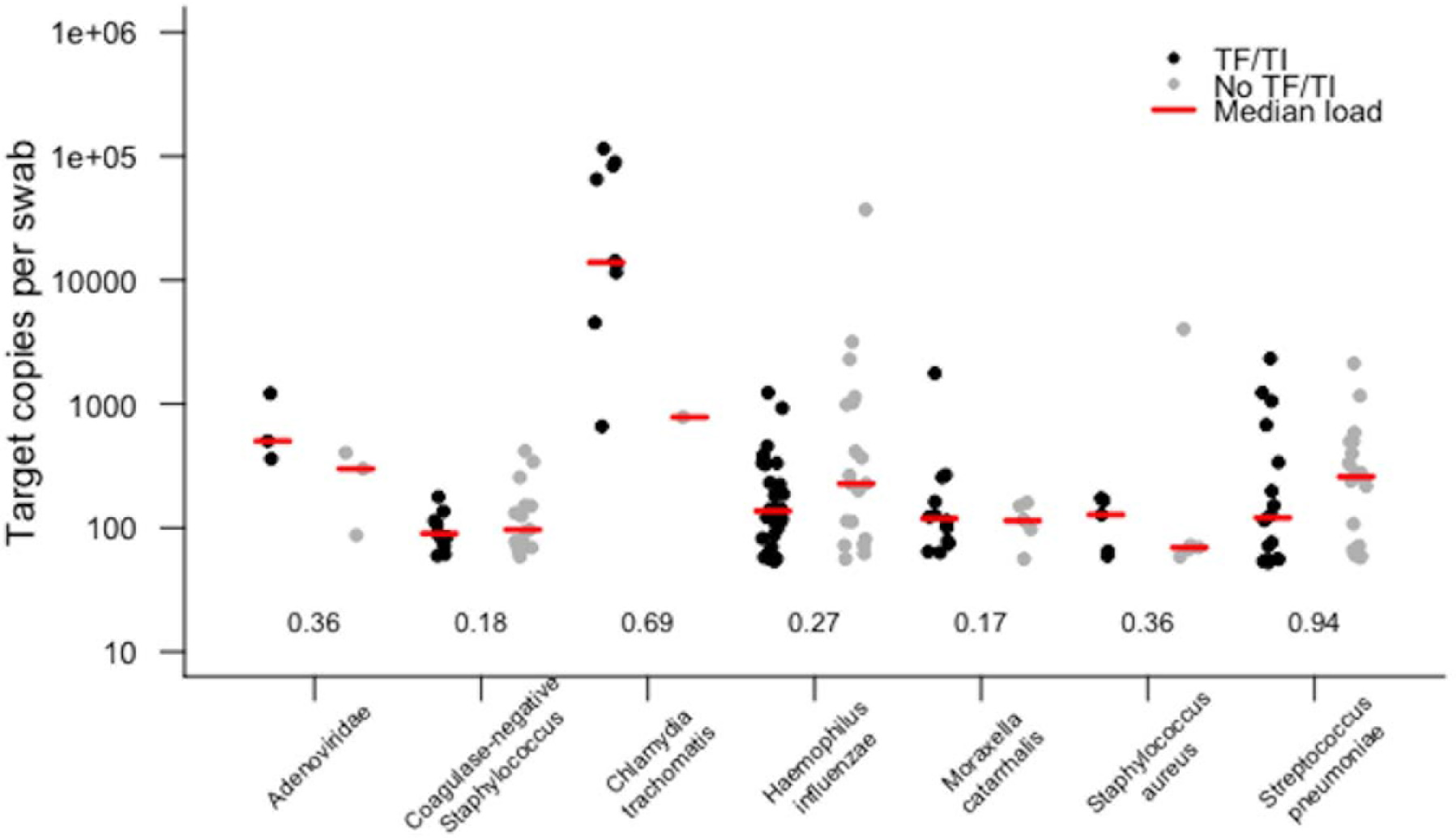
Target copies per swab of each pathogen identified in conjunctival swabs collected from children with and without active trachoma in the Solomon Islands. Numbers show p-values for logistic regression comparison between active trachoma case and control groups for each pathogen. None of the differences observed were statistically significant.

### 16S rRNA gene sequencing

The total number of genera identified across all clinical specimens and NTCs was 659, however, 125/659 (19%) had a cumulative total of ten reads or fewer. The median number of reads per clinical specimen was 55,104 (IQR: 40,705 − 89,541) and the median percentage of reads mapped to genus level was 51.9%. The median number of genera identified was 151 (IQR: 117 − 212) per clinical specimen. Following removal of presumed contaminants, approximately 40% of cleaned reads were assigned to genera that each constituted less than 1% of the total polymicrobial community. 40 genera were represented by at least 1% of remaining reads in clinical samples.

The diversity of bacterial communities was in general low; the mean Inverse Simpsons Diversity index over all specimens was 0.061 (IQR: 0.046 − 0.092). The median Inverse Simpson’s Diversity index in cases was 0.061 [IQR: 0.048 − 0.095] versus controls 0.060 [IQR: 0.044 − 0.078] (Student’s t-test p = 0.56). The median Shannon Diversity Index was 3.38 in cases and 3.37 in controls (Student’s t-test p = 0.96). There was no significant difference in alpha-diversity between cases and controls by either measure of diversity. In cases of active trachoma, the most dominant bacterial genera were *Corynebacterium* (12.0% of total reads), *Proprionibacterium* (6.2%) and *Helicobacter* (4.8%). In controls, the most dominant genera were *Corynebacterium* (13.9%), *Paracoccus* (5.2%), *Proprionibacterium* (4.7%), and *Neisseria* (4.1%) (Supplementary Figure 1). The genus level membership of the bacterial community varied between cases and controls. Of 21 genera found in specimens from those with active trachoma, 10 were not found in controls, including *Helicobacter*, *Mesoplasma*, *Brachybacterium* and *Haemophilus*. Of 23 genera found in controls, 12 were not found in cases, including *Neisseria*, *Prevotella*, *Rhodococcus* and *Porphyromonas*.

Principal Components (PC) Analysis (Figure 2, Supplementary Figure 2) revealed that *Corynebacterium* (PC1), *Paracoccus* (PC2), *Propionibacterium* (PC2, PC3) and *Helicobacter* (PC3) are major contributors to the variation in the conjunctival microbiomes of children in the Solomon Islands. While individual cases or controls have distinctive profiles dominated by these genera, the majority of cases and controls are indistinguishable (Figure 2) using the first three PCs (explaining 45% of total variation). We condensed PCs 1-20 into a single discriminant function, which explained 87% of variation in the data. In this analysis, the genus-level polymicrobial community structure is significantly different between cases and controls (logistic regression p = 0.0000062) (Figure 3). That discriminant function was dominated by a number of genera such as *Curvibacterium* and *Mesoplasma* (Supplementary Figure 3). Cross validation to predict group membership using discriminant functions of between 10 and 90 PCs had a low success rate (median 53.3% success, range: 50.1 − 62.3%). This suggests that while differences do exist between the polymicrobial communities in cases and controls, they are too subtle and varied to be predictive of phenotype, at least when working with this number of specimens.

**Figure 2.**
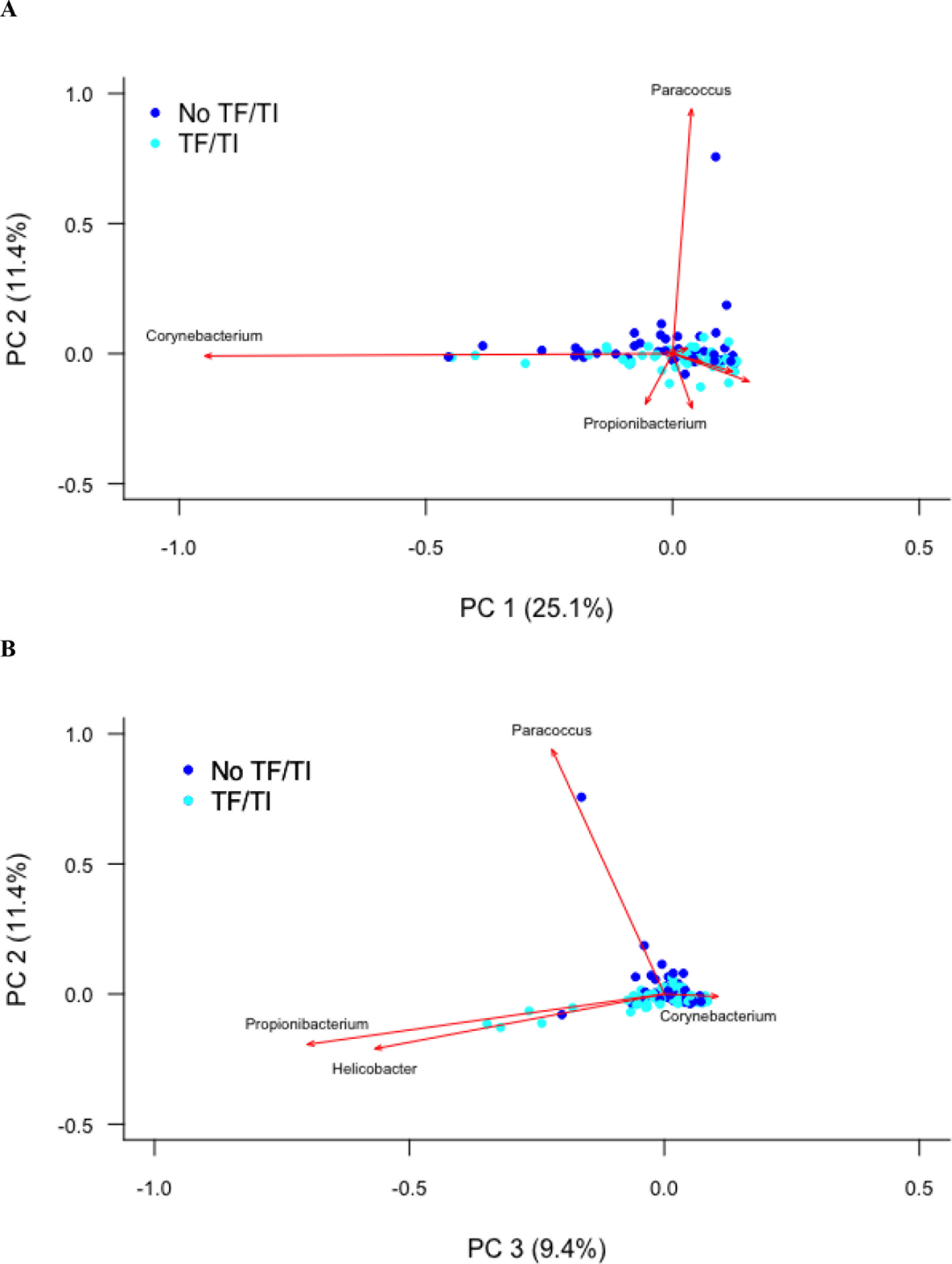
**(A)** First and second and **(B)** second and third principal components describing variation between the 16S sequences identified in Solomon Island children with and without active trachoma. Dark blue spots indicate controls. Light blue spots indicate cases. Red arrows show principal component loadings.

**Figure 3:**
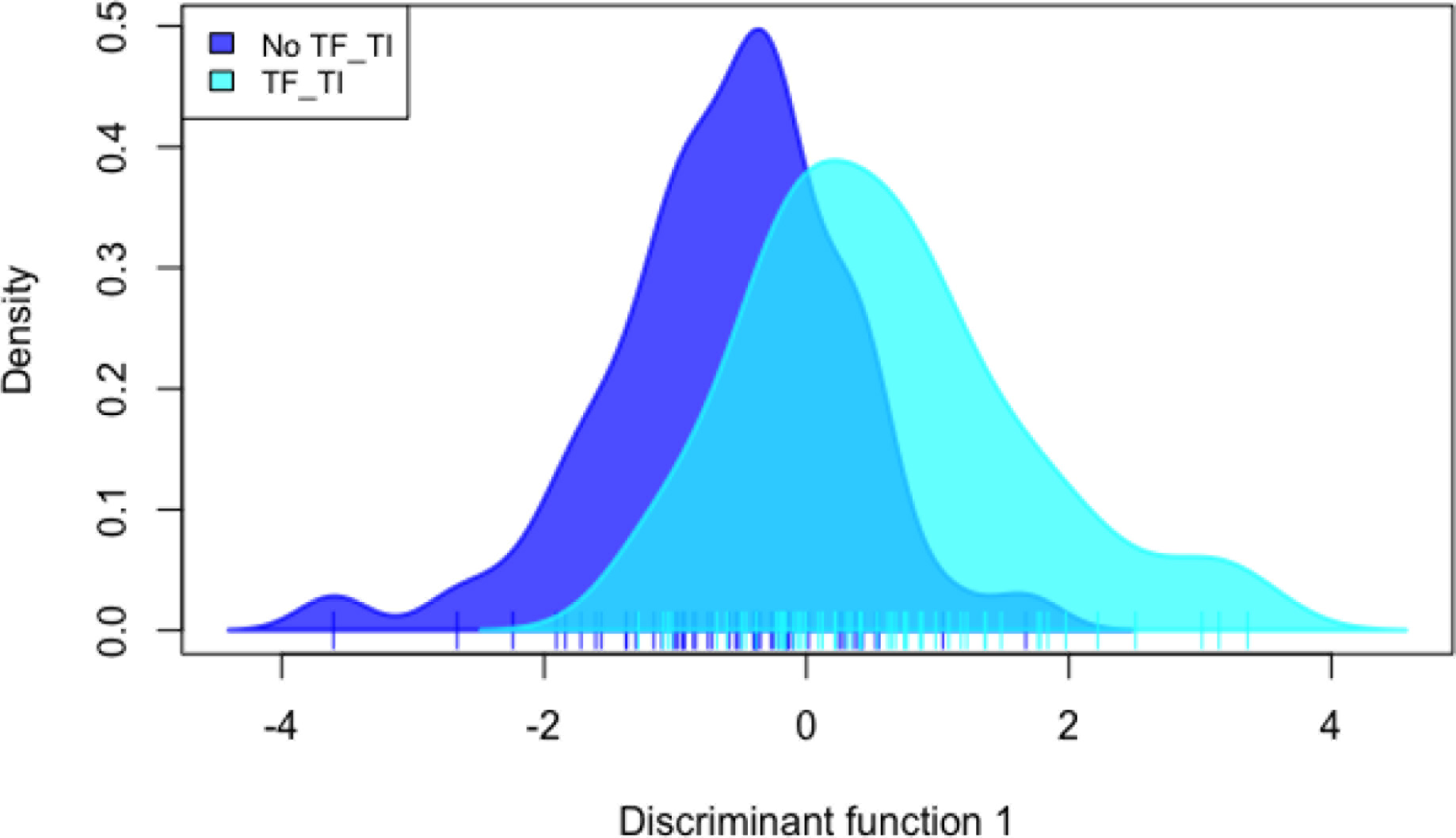
Discriminant analysis of the association of 20 combined principal components with active trachoma, showing a significant discrimination of phenotype groups (p = 0.000006).

## Discussion

Follicular conjunctivitis meeting criteria for the trachoma phenotype TF is highly prevalent in the Solomon Islands, but the prevalence of *Ct* infection is curiously low. Although *Ct* was the only pathogen to associate with TF in this study, the prevalence was much lower than may be expected of a population with this level of TF (3). Both *S. pneumoniae* and *H. influenzae* were detected at moderate prevalence in our population, whilst we found very few cases of *Adenoviridae* and *M. catarrhalis* infection. Comparator data from other populations are scarce, but it is clear that staphylococcal species were detected in the conjunctivae in the Solomon Islands at a substantially lower prevalence (7%) than in children in either Tanzania (14.8% *S. epidermidis*) (7), Sierra Leone (20% *S. aureus*, 29% coagulase-negative *Staphylococcus*) (38) or The Gambia (post MDA, 14.7% *S. aureus*) (11). By ddPCR testing for a number of bacteria and viruses that have previously been linked to TF, we can discount the possibility that any of them can account either singly or *en masse* for the high TF prevalence.

Through 16S amplicon sequencing and community profiling, we have shown that there is apparently no dominant bacterial genus associated with this disease. The dominant features of variation in conjunctival bacterial communities in Solomon Islands children (*Corynebacterium*, *Propionibacterium*) have consistently been identified in other studies. Other genera (e.g., *Paracoccus*, *Helicobacter*, *Haemophilus*) have not previously been identified in 16S studies, whilst some reported in other studies (e.g., *Simonsella*, *Pseudomonas*) were not found in these specimens (19–22). Many studies have sequenced the V3-4 region of the 16S gene whereas we targeted V1-3. V region choice has been shown to have an impact on the genera identified at other mucosal sites (39) and, while no data have yet emerged on how this affects profiles from the eye, it is possible this could account for some of the differences. Consistent with previous studies (36), we found a background of genera that amplified in NTCs that also appeared in clinical specimens. The microbiological biomass at the conjunctiva is known to be very low, even compared with other nearby sites such as skin or oral mucosa (20). When true resident bacteria are scarce, 16S rRNA gene PCR readily amplifies reagent contaminants. We took stringent measures to exclude reagent contaminants, but this resulted in the removal of some important genera such as *Staphylococcus* and *Streptococcus*. Among the biggest contributors to between-conjunctiva variation were *Corynebacterium* and *Propionibacterium*, but these genera did not differentiate TF cases from controls (Figure 2). Previous investigations of the role of the conjunctival microbiome in ocular disease have also not shown significant differences. In The Gambia, there was an increased abundance of *Haemophilus* in cases of TF, as compared to controls, although this difference was not significant (22). People with ocular manifestations of rosacea, Sjögren’s syndrome and healthy controls did not have significant differences in diversity or relative abundance of key phyla identified across all three groups (24). However, other disease associations have been identified. For example, the abundance of *Corynebacterium* and *Streptococcus* varied significantly between individuals with and without conjunctival scarring (22).

The microbiota of these children were heterogeneous and had subtle variations that appeared to reach significant association with TF when the full community structure was considered in comparative statistical models. It remains possible (though perhaps unlikely) that a multitude of factors, operating at the level of single bacterial species, polymicrobial communities or viral species, all contribute to the presentation of the phenotype. It is perhaps more likely that as yet unidentified viral or allergic causes could explain the prevalence of TF in the Solomon Islands.

There are some limitations to our study. Firstly, although the ddPCR assays are based on validated assays and appear to be accurate (Supplementary Table 1), they have not been formally evaluated in this format. Secondly, by not using mechanical lysis in our extraction process, the absolute prevalence of some difficult-to-lyse Gram positive genera (e.g., *Staphylococcus*) may have been under-estimated. This would not impair the comparison of cases to controls, between which protocols were consistent. Thirdly, 16S amplicon sequencing of low biomass samples is known to result in amplification of reagent and environmental contaminants (36). This study focused on gross differences in community structure between those with and without disease and these should be independent of contaminating reads. We also employed stringent quality control measures to ensure data were not confused by artefacts of the sequencing process.

Given the current international commitment to trachoma elimination, further characterisation of the role of non-*Ct* stimuli in TF is important. We might expect those with TF in the absence of *Ct* to be at lower risk of progression to TS and TT than those with repeated *Ct* infection and inflammation (40), however, such individuals have not been studied longitudinally to assess outcomes therefore it is unclear how these cases should be managed. Furthermore, years after MDA has been administered, TF persists at levels above the threshold for elimination (>5%) in communities where *Ct* infection is not readily detectable (41, 42). The public health risk of trachoma in these communities is also unclear.

Public health scenarios such as the one in the Solomon Islands will become increasingly common as trachoma elimination programmes reduce the global prevalence of *Ct*, and the positive predictive value of TF for ocular Ct infection declines as a result (43). MDA might be inappropriately delivered when clinical signs of trachoma are the only indicator used for programmatic decision making. While MDA is effective for the treatment of trachoma (44), mass antibiotic exposure may increase macrolide resistance in nonchlamydial bacteria (45) and theoretically could make the population more susceptible to later infections by preventing accumulation of acquired immunity to *Ct* (46). Decisions to undertake such a program should therefore be carefully considered. The case for using tests for *Ct* infection during trachoma surveys is strengthened by our data. The value of nucleic acid amplification tests for detecting non-chlamydial infection remains questionable, as multiple infectious agents might be important, and these are likely to differ between populations.

## Acknowledgements

We thank Victoria Miari and Emma Cobb (Pathogen Molecular Biology Department, London School of Hygiene & Tropical Medicine, UK) for providing bacterial culture isolates for PCR assay validation; and David Nelson (Indiana University, Indianapolis, US) for the provision of 16S sequencing primers and protocols. We would also like to thank the residents of the communities who took part in this study.

## Financial statement

Fieldwork was jointly funded by the United Kingdom’s Department for International Development Global Trachoma Mapping Project grant (ARIES: 203145) to Sightsavers, and by the Fred Hollows Foundation, Australia (1041). Laboratory costs were funded by the Fred Hollows Foundation, Australia (1041).

RMRB and AWS were funded by a Wellcome Trust Intermediate Fellowship to AWS (098521).

OS, KJ, EK, LS and CR were seconded by the Solomon Islands Ministry of Health and Medical Services for the duration of the survey.

MJH is supported by the Wellcome Trust (093368/Z/10/Z).

ChR is supported by the Wellcome Trust Institutional Strategic Support Fund (105609/Z/14/Z).

## Author Contributions

Conceived and designed the study: RMRB, OS, RTLM, AWS, DCWM, ChR.

Performed the fieldwork: RMRB, OS, KJ, EK, LS, CR.

Performed the experiments: RMRB, MJH, JH, CP, ChR.

Analysed the data: RMRB, ChR.

Wrote the manuscript: RMRB, ChR.

Revised and approved the manuscript: RMRB, OS, KJ, EK, LS, CR, JH, CP, MJH, RTLM, AWS, DCWM, ChR.

